# Enthalpy efficiency of the soleus muscle explains improvements in running economy

**DOI:** 10.1101/2020.07.09.194654

**Authors:** Sebastian Bohm, Falk Mersmann, Alessandro Santuz, Adamantios Arampatzis

**Affiliations:** Humboldt-Universität zu Berlin, Department of Training and Movement Sciences, Philippstr. 13, 10115 Berlin, Germany; Berlin School of Movement Science, Humboldt-Universität zu Berlin, Berlin, Germany

## Abstract

During human running, the soleus, as the main plantar flexor muscle, generates the majority of the mechanical work through active shortening. The fraction of chemical energy that is converted into muscular work (i.e. the enthalpy efficiency) depends on the muscle shortening velocity. Here, we investigated the soleus muscle fascicle behavior during running with respect to the enthalpy efficiency as a mechanism that could explain previously reported improvements in running economy after exercise-induced increases of plantar flexor strength and Achilles tendon stiffness. Healthy amateur runners were randomly assigned to a control (n=10) or intervention group (n=13), which performed a specific 14-week muscle-tendon training. Significant increases in plantar flexor maximum strength (10%) and Achilles tendon stiffness (31%) yet reduced metabolic cost of running (4%) was found only in the intervention group (p<0.05). Following training, the soleus fascicle velocity profile throughout stance was altered, with the fascicles operating at a higher enthalpy efficiency during the phase of muscle-tendon unit lengthening (15%) and in average over stance (7%, p<0.05). These findings show that the improvements in energetic cost following increases in plantar flexor strength and Achilles tendon stiffness can be attributed to increased enthalpy efficiency of the operating soleus. This provides the first experimental evidence that the soleus enthalpy efficiency is a determinant of human running economy. Furthermore, the current results imply that the soleus energy production in the first part of the stance phase were the muscle-tendon unit is lengthening is crucial for the overall metabolic energy cost of running.

## Introduction

Habitual bipedalism has been recognized as a defining feature of humans [1] and an exceptional endurance running ability has been linked to the evolution of the human lineage [2–4]. Economy, which is the mass-specific rate of oxygen uptake or metabolic energy consumption at a given speed [5,6], plays a crucial role for endurance running performance [7]. The cost of generating force and work through muscles to support and accelerate the body mass is the main source of metabolic energy expenditure during locomotion [8,9]. The force-length-velocity potential of muscles, defined as the fraction of maximum force according to the force-length [10] and force-velocity relationships [11], at which muscles operate during running [12,13] largely dictates the required active muscle volume and consequently the energetic cost of contraction [5,12,14].

In human running, the triceps surae is the major contributor to propulsion and the main plantar flexor muscle group that transmits force through the Achilles tendon (AT) [15,16], consuming a significant amount of metabolic energy [17]. In earlier studies, we provided evidence that both the contractile capacities of the triceps surae and the mechanical properties of the AT, i.e. its stiffness, influence running economy [18,19]. We found that the most economical runners feature a combination of higher plantar flexor muscle strength and AT stiffness [18] and that a specific training of muscle strength and AT stiffness can in fact improve running economy [19]. Although the association of the AT stiffness and energetic cost of running has been confirmed by other research groups [20,21], the underlying physiological mechanisms are yet unclear.

The soleus is the greatest muscle of the triceps surae [22] and generates the majority of work/energy to lift and accelerate the body [15] by actively shortening throughout the entire stance phase of running [12,23]. In the first part of the stance, the fascicle shortening is paralleled by a lengthening of the muscle-tendon unit (MTU) [12], indicating that a part of the body’s mechanical energy is stored as strain energy in the AT, but also that the fascicles generate work and save this work as strain energy in the AT. In the second part of the stance phase, where the MTU shortens (propulsion phase), the tendon strain energy is returned to the body and contributes to the ongoing work generation by active fascicle shortening [12]. The metabolic cost of generating work by active shortening of muscles depends on the velocity of the shortening [24]. The enthalpy efficiency (or mechanical efficiency) quantifies the fraction of chemical energy from ATP hydrolysis that is converted into mechanical muscular work [25]. The relation of enthalpy efficiency and shortening velocity shows a steep increase at low velocities with the peak at around 20% of the maximum shortening velocity [25,26]. During submaximal running, the soleus operates below the optimal velocity for maximal efficiency [12], suggesting that small changes in the shortening velocity may substantially influence the enthalpy efficiency of the soleus muscular work production.

The mechanical interaction of the soleus muscle with the series AT regulates the fascicle shortening dynamics. The AT takes over a great part of the length changes of the entire soleus MTU, thereby decoupling the muscle fascicle and MTU behavior and, besides the storage and release of strain energy, allowing the fascicles to operate at velocities favorable for economical force generation [12,23]. The mechanical properties of the tendon in combination with the strength capacity of the muscle may determine the amount of fascicle decoupling during the stance phase of running. However, similar to an increase in muscle strength [27], tendons can adapt to periods of higher mechanical loading by increasing stiffness [28,29]. Our earlier findings of improved energetic cost after an exercise-induced increase in AT stiffness and muscle strength evidenced a direct association between a balanced adaptation of tendon and muscle and improvements in running economy [19]. Considering a given work produced by the soleus muscle during the stance phase of running, the energetic cost depends on the enthalpy efficiency under which this muscular work is generated. Assuming that a combination of increased plantar flexor strength and AT stiffness may influence the soleus fascicle shortening pattern during the stance phase of running, the overall enthalpy efficiency might improve. This would provide an explaining mechanism to the previously reported improvements in running economy following an effective muscle-tendon training [19]. To the best of our knowledge, no study experimentally examined the operating soleus muscle fascicles with respect to the enthalpy efficiency and its association to the energetic cost of running.

Here, we investigated the effect of a specific muscle-tendon training, which has been shown to increase muscle strength of the plantar flexors as well as AT stiffness [19], on the enthalpy efficiency of the operating soleus fascicles during running. Based on our earlier study [19], we expected an improvement in running economy after 14 weeks of training. We hypothesized that the training-induced increase in muscle strength and AT stiffness modulates the soleus fascicle velocity pattern throughout the stance phase towards velocities associated with a higher enthalpy efficiency of the operating soleus muscle, thereby reducing the energetic cost of running.

## Methods

### Participants and experimental design

A statistical power analysis was performed *a priori* to calculate the required sample size by means of the software G*Power (version 3.1.9.6, HHU Düsseldorf, Germany) [30]. For this purpose, we used the effect size of the rate of oxygen consumption from our previous intervention study with the same training regimen (d = 1.04) [19]. Since the main outcome of interest was the effect of training, the power analysis was conducted for the *post-hoc* time point comparison for the intervention group considering a Bonferroni correction of the p-values (α = 0.025 (adjusted), power 0.8, two-tailed paired t-test). The analysis revealed a required sample size of n = 12 for the intervention group. Under consideration of potential dropouts, we recruited 36 participants and randomly assigned them to either an intervention (n = 19) or control group (n = 17). Inclusion criteria were age 20 to 40 years, at least three times per week running training and no severe muscular-tendinous injuries in the previous year. Only habitual rearfoot-striking runners were considered. To quantify the foot strike pattern, we assessed the strike index [31], a measure of the position of the center of pressure with respect to the heel relative to foot length at touchdown, during a pretest session. A strike index of 0 indicates extreme rearfoot striking and of 1 extreme forefoot striking, while <0.3 was set as the threshold for the study inclusion. Due to injuries (not related to the intervention) and time required for the assessment and/or training, 13 participants canceled their commitment. Twenty-three participants completed the intervention, 13 in the intervention group (age: 29 ± 5 years, height: 178 ± 8 cm, mass: 73 ± 8 kg, 4 female) and 10 in the control group (age: 31 ± 3 years, height: 175 ± 10 cm, mass: 70 ± 11 kg, 7 female). For the intervention group, the same 14-week muscle-tendon training was added to the regular ongoing training habits as in our earlier study [19]. Before and after the intervention period, the maximal plantar flexion moment and AT stiffness as well as energetic cost of running at 2.5 m/s were assessed in both groups. In order to explain the expected improvements in running economy (i.e. energetic cost) following the muscle-tendon training, we experimentally determined a) the foot strike pattern and temporal gait parameters as well as b) the soleus MTU and fascicle behavior in addition to the soleus electromyographic (EMG) activity during running. We further determined c) the soleus force-fascicle length relationship and force-fascicle velocity relationship in order to calculate the force-length and force-velocity potential of the fascicles during running (i.e. fraction of maximum force according to the force-length and force-velocity curve [12,13,32]) and assessed d) the efficiency-fascicle velocity relationship to calculate the efficiency of the soleus during running. Because changes in running economy were not expected without any intervention [19], the assessment of the soleus fascicle behavior was not conducted in the control group. The ethics committee of the Humboldt-Universität zu Berlin approved the study and the participants gave written informed consent in accordance with the Declaration of Helsinki.

### Exercise protocol

The supervised resistance training program was performed for 14 weeks and was charactarized by five sets per session of four repetitive isometric ankle plantarflexion contractions (3 s loading, 3 s relaxation) at 90% of the maximum voluntary contraction (MVC) strength (adjusted every two weeks), three to four times a week. The ankle joint was set to 5° dorsiflexion and the knee joint was fully extended. This loading regimen has been shown to provide a sufficient magnitude and duration of strain to promote AT adaptation in addition to increases in muscle strength of the plantar flexors [19,29,33]. Online biofeedback of the strength effort was displayed to control the desired loading.

### Strength of the plantar flexors and Achilles tendon stiffness

The strength of the plantar flexors of the right leg was measured before and after the 14 weeks in a seated position (70° of hip flexion) with the knee extended using a Biodex dynamometer (Biodex Medical Inc., Syst.3, Shirley, NY, USA). Following a standardized warm-up, five MVCs were performed in resting ankle joint angles of 10° dorsiflexion to the individual maximum dorsiflexion angle (0° = foot perpendicular to shank) in equally distributed intervals (3-7°) to determine the maximum joint moment. The resultant ankle joint moment was calculated using an established inverse dynamics approach to account for misalignments between dynamometer and joint axis as well as passive and gravitational moments [34,35]. Furthermore, the contribution of the antagonistic muscles to the ankle joint moment was considered by means of an EMG-based method [36].

For the determination of AT stiffness, five ramp-MVCs with steadily increasing effort from rest to maximum under the same considerations (i.e. accounting for axis misalignment, passive and gravitational moments and co-activation) were conducted at 0° ankle angle. The force applied to the AT was calculated as quotient of the joint moment and the individual tendon lever arm, which was determined using the tendon-excursion method [37,38] and corrected for tendon alignment during the contraction [39]. The corresponding AT elongation during the ramp MVCs was analyzed based on the displacement of the gastrocnemius medialis-myotendinous junction (MTJ) visualized by B-mode ultrasonography captures (My Lab 60, Esaote, Genova, Italy, 25 Hz). The MTJ displacement artefacts due to an unavoidable change in the ankle joint angle during the MVCs was corrected [40] and the five contractions were averaged to give a reliable measure of the elongation [41]. The AT stiffness was calculated between 50% and 100% of the maximum tendon force using linear regression [29]. In order to calculate AT strain, the rest length was measured from the tuberositas calcanei to the MTJ at an ankle angle of 110° (plantar flexed) and extended knee (i.e. a position that provides AT slackness [42]).

### Energetic cost of running

During a 10-minute running trial on a treadmill (h/p cosmos mercury, Isny, Germany) at 2.5 m/s, a breath-by-breath cardio pulmonary exercise testing system (MetaLyzer 3B-R2, CORTEX Biophysik GmbH, Leipzig, Germany) recorded the percentage of concentration of oxygen and carbon dioxide expired. Rate of oxygen consumption 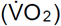 and carbon dioxide production 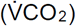 was calculated as average of the last three minutes [43]. Running economy was expressed in units of energy by:

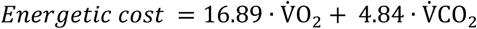

where the energetic cost is presented in [W/kg] and 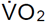 and 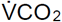 in [ml/s/kg] [6,44]. To reduce test-retest variability [45], the shoes, time of testing and training activity during the previous 72 hours were the same for the pre and post measurements. No food intake was allowed during the last 3 hours before testing.

### Joint kinematics and foot strike pattern

During running, kinematics of the right leg were captured by a Vicon motion capture system (250 Hz) using anatomical-referenced reflective markers (greater trochanter, lateral femoral epicondyle and malleolus, fifth metatarsal and calcaneus). The touchdown of the foot and the toe-off was determined from the kinematic data as consecutive minimum in knee joint angle over time [46].

The foot strike pattern was analyzed by means of the strike index [31] determined during the ten-minute running trial. A self-developed algorithm [47] was used to calculate the strike index from the plantar pressure distribution (120 Hz) captured by the integrated pressure plate (FDM-THM-S, Zebris Medical GmbH, Isny, Germany).

### Soleus muscle-tendon unit length changes, fascicle behavior and electromyographic activity during running

During an additional 3-minute running trial at the same speed, kinematics of the ankle joint served to calculate the length change of the soleus MTU as the product of ankle angle changes and the previously assessed individual AT lever arm [48]. The initial soleus MTU length was determined at neutral ankle joint angle based on the regression equation provided by Hawkins & Hull [49]. Ultrasonic images of the soleus muscle fascicles were obtained synchronously to the kinematic data at 146 Hz (Aloka Prosound Alpha 7, Hitachi, Tokyo, Japan). The ultrasound probe (6 cm linear array, UST-5713T, 13.3 MHz) was mounted on the shank over the medial aspect of the soleus muscle belly using a custom antiskid neoprene-plastic cast. The fascicle length was post-processed from the ultrasound images using a self-developed semi-automatic tracking algorithm [50] and corrections were made if necessary during visual inspection of each image. At least nine steps were analyzed for each participant and averaged [13,51]. The velocities of MTU and fascicles were calculated as the first derivative of the lengths over the time. Synchronized surface EMG of soleus was measured (1000 Hz) by means of a wireless EMG system (Myon m320RX, Myon AG, Baar, Switzerland). The EMG data are presented as normalized to the maximum EMG value observed from the individual MVCs [12].

### Soleus force-length, force velocity and efficiency-velocity relationship

To determine the soleus force-fascicle length relationship (for details see [12]), the participants were placed in prone position on the bench of the dynamometer with the knee fixed in flexed position (fig. 1) to restrict the contribution of the bi-articular muscle gastrocnemius to the plantar flexion moment (∼120°) [52]. MVCs were performed with the right leg in eight different joint angles, equally distributed between 10° plantar flexion to the individual maximum dorsiflexion and executed in a randomized order. The joint moments were calculated using the approach described above. The force acting on the AT was calculated by dividing the joint moment by the previously determined AT lever arm. The corresponding soleus fascicle behavior during the MVCs was captured synchronously at 30 Hz by ultrasonography and fascicle length was measured as described above (fig. 1). The ultrasound probe remained attached between the running trial and MVCs. An individual force-fascicle length relationship was calculated by means of a second-order polynomial fit (fig. 1), giving the individual maximum force and optimal fascicle length for force generation (L_0_).

**Fig. 1:**
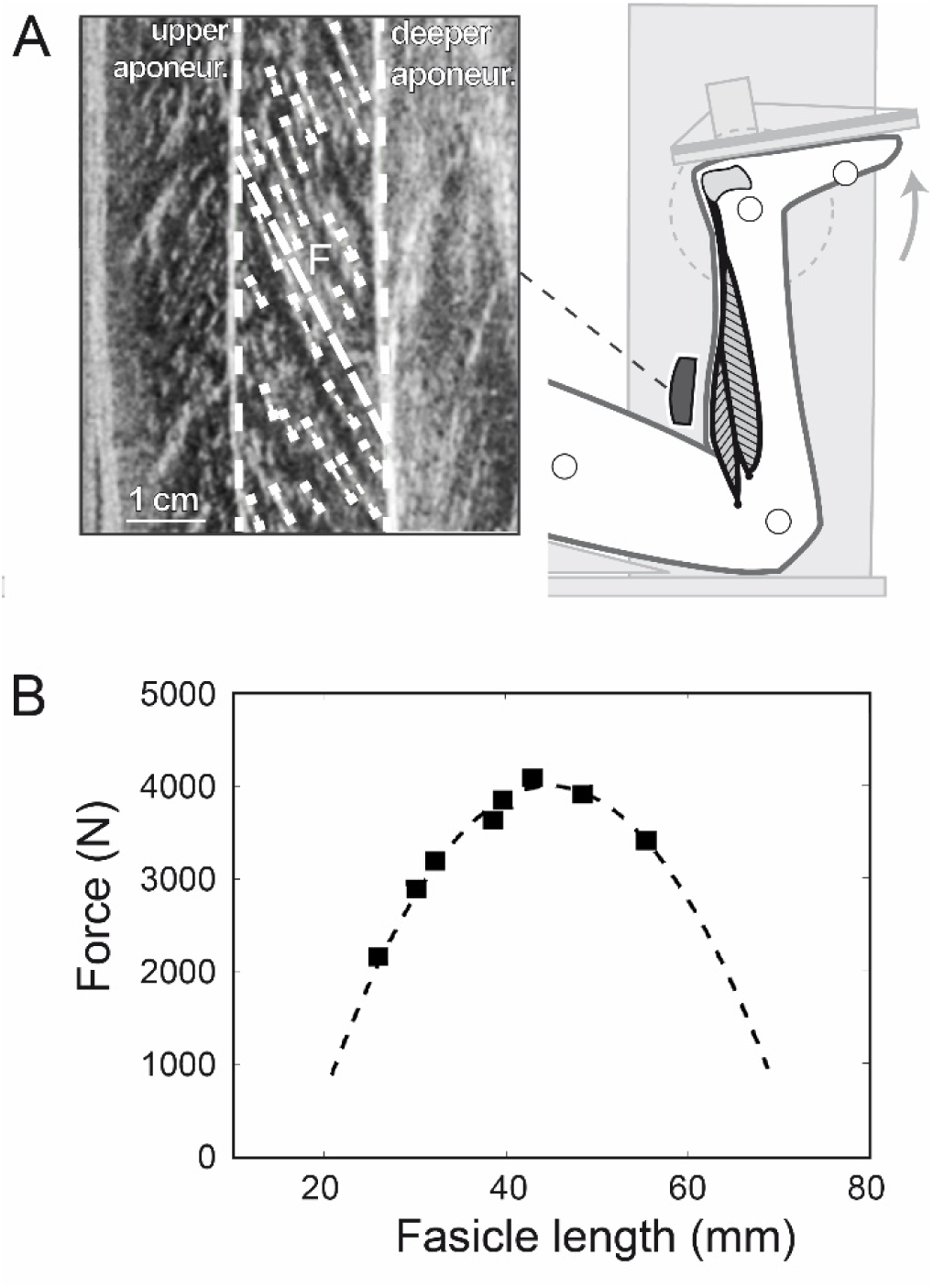
Experimental setup for the determination of the soleus force-fascicle length relationship. A) Maximum isometric plantar flexions (MVCs) at eight different joint angles were performed on a dynamometer. During the MVCs, the soleus muscle fascicle length was measured by ultrasonography as an average (*F*) of multiple fascicle portions (short dashed white lines) identified from the images. B) Exemplary force-length relationship of the soleus fascicles obtained from the MVCs (squares) and the respective second-order polynomial fit (dashed line).

The force-velocity relationship of the soleus was assessed using the classical Hill equation [11] and the muscle-specific maximum fascicle shortening velocity (V_max_) and constants of a_rel_ and b_rel_. For V_max_ we took values of human soleus type 1 and 2 fibers measured in vitro at 15°C reported by Luden et al. [53]. The values were then adjusted for physiological temperature conditions (37 °C) using the temperature coefficients provided by Ranatunga [54]. An average fiber type distribution (type 1 fibers: 81%, type 2: 19%) of the human soleus muscle reported in literature [53,55–57] was then the basis to calculate a representative value of V_max_ of the soleus in vivo as 6.77 L_0_ /s [12], were L_0_ refers to the individual measured optimal fascicle length. a_rel_ was calculated as 0.1 + 0.4FT, where FT is the fast twitch fiber type percentage (see above), which then equals to 0.175 [58,59]. The product of a_rel_ and V_max_ gives b_rel_ as 1.182 [60]. Based on the assessed force-length and force-velocity relationship, it was possible to calculate the individual force-length and force-velocity potential of the soleus muscle as a function of the fascicle operating length (fig. 1) and velocity during running (i.e. force-length and force-velocity potential as fraction of maximum force according to the force-length and force-velocity curve [12,13,32]).

Furthermore, we determined the enthalpy efficiency-shortening velocity relationship for the soleus muscle fascicles to calculate the enthalpy efficiency of the soleus muscle as a function of the fascicle operating velocity during running. For this purpose, we referred to the experimental efficiency values provided by the paper of Hill 1964 in table 1 [24]. The original values were presented as a function of relative load (relative to maximum tension) which we then transposed to the shortening velocity (normalized to maximum shortening velocity) on the basis of the classical Hill equation [11]. The corresponding values of enthalpy efficiency and shortening velocity were then fitted using a cubic spline, giving the right-skewed parabolic-shaped curve with a peak efficiency of 0.45 at a velocity of 0.18 V_max_. The resulting function was then used to calculate the efficiency of soleus during running.

**Table 1:** Maximal plantar flexion moment and Achilles tendon stiffness as well as energetic cost, foot strike index and temporal step characteristics during running before and after the training period for the intervention and control group (mean±SD, effect size g).

|  | Intervention (n = 13) |  |  | Control (n = 10) |  |  |
| --- | --- | --- | --- | --- | --- | --- |
|  | Pre | Post | g | Pre | Post | g |
| Moment (Nm/kg)<sup>+</sup> | 3.12 ± 0.48 | 3.44 ± 0.37* | 0.77 | 3.10 ± 0.46 | 2.99 ± 0.32 | 0.32 |
| Stiffness (kN/strain)<sup>+</sup> | 85 ± 36 | 111 ± 59* | 0.67 | 73 ± 29 | 71 ± 28 | 0.10 |
| Energy cost (W/kg)<sup>#</sup> | 10.6 ± 0.6 | 10.2 ± 0.7* | 0.74 | 11.2 ± 1.0 | 11.1 ± 1.0 | 0.12 |
| Strike index | 0.08 ± 0.12 | 0.10 ± 0.16 | 0.09 | 0.06 ± 0.03 | 0.06 ± 0.03 | 0.05 |
| Stance time (ms) | 310 ± 23 | 316 ± 23 | 0.29 | 327 ± 17 | 324 ± 23 | 0.34 |
| Flight time (ms) | 53 ± 31 | 53 ± 24 | 0.01 | 50 ± 31 | 54 ± 31 | 0.48 |
| Cadence (steps/min) | 160 ± 11 | 159 ± 9 | 0.39 | 162 ± 9 | 161 ± 9 | 0.26 |

<sup>+</sup> Significant time by group interaction effect (p < 0.05)
<sup>#</sup> Significant main effect of time (p < 0.05)
\* Significant difference (post hoc) to pre (p < 0.05)

**Table 2:**
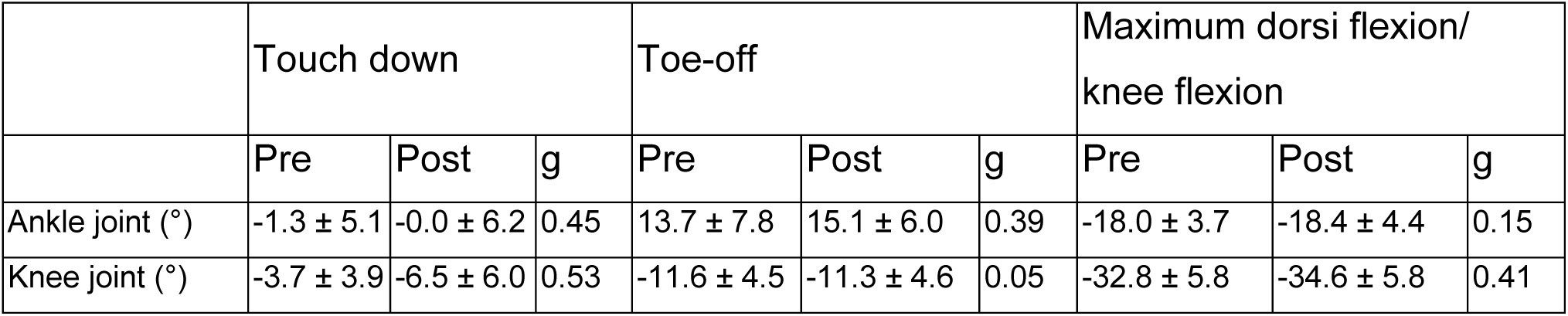
Ankle and knee joint angles at touch down, toe-off and at the maximal ankle dorsiflexion and knee flexion angle, respectively, during running before and after the training intervention (mean±SD, effect size g, n=13).

**Table 3:** Soleus muscle-tendon unit (MTU) length, length changes and velocity as well as muscle fascicle length, fascicle shortening distance and fascicle velocity averaged over the phase of MTU lengthening, MTU shortening and over the entire stance phase during running before and after the training intervention (mean±SD, effect size g, n=13).

|  | MTU lengthening |  |  | MTU shortening |  |  | Stance phase |  |  |
| --- | --- | --- | --- | --- | --- | --- | --- | --- | --- |
|  | Pre | Post | g | Pre | Post | g | Pre | Post | g |
| MTU length (mm) | 325 ± 20 | 325 ± 21 | 0.02 | 323 ± 20 | 323 ± 21 | 0.02 | 324 ± 20 | 324 ± 21 | 0.02 |
| MTU length changes (mm) | 18.4 ± 2.0 | 18.2 ± 3.2 | 0.04 | 33.9 ± 9.3 | 32.5 ± 5.6 | 0.19 | 16.4 ± 9.0 | 14.8 ± 5.6 | 0.30 |
| MTU velocity (mm/s) | -97 ± 15 | -98 ± 22 | 0.07 | 259 ± 52 | 244 ± 33 | 0.36 | 173 ± 29 | 164 ± 21 | 0.30 |
| Fasicle length (mm) | 39.2 ± 4.4 | 39.0 ± 5.1 | 0.03 | 34.5 ± 4.3 | 33.1 ± 4.5 | 0.23 | 37.2 ± 4.3 | 36.5 ± 4.8 | 0.12 |
| Fascicle shortening (mm) | 5.21 ± 2.68 | 6.75 ± 3.08 | 0.45 | 6.49 ± 2.02 | 4.98 ± 1.23* | 0.72 | 11.05 ± 3.32 | 11.53 ± 3.47 | 0.12 |
| Fasicle velocity (mm/s) | 21.2 ± 16.7 | 33.4 ± 17.5 | 0.52 | 49.1 ± 16.7 | 35.8 ± 10.1* | 0.71 | 33.0 ± 10.8 | 34.6 ± 11.0 | 0.10 |

\* Significant difference to pre (p < 0.05)

### Statistics

An analysis of variance for repeated measures was performed for the plantar flexion moment (normalized to body mass) and AT stiffness (normalized to resting length) as well as metabolic energy cost, foot strike index and temporal gait characteristics during running with the time point as the within-subjects factor (pre vs. post) and the group as a between-subjects factor (intervention vs control). The *post-hoc* analysis was conducted separately for each group considering a Benjamini-Hochberg correction (adjusted p-values reported). Normality of the standardized residuals was controlled using the Kolmogorov-Smirnov test with Lilliefors correction.

Anthropometric group differences as well as baseline differences of the plantar flexion moment, AT stiffness and energetic cost were tested using a t-test for independent samples. A paired t-test was used to analyze the training effects on the assessed gait characteristics, kinematics and MTU and fascicle parameters. If normality tested by the Kolmogorov-Smirnov test was not given, the Wilcoxon signed rank test was applied. The level of significance was set to α = 0.05 and the statistical analyses were performed using SPSS (IBM Corp., version 22, NY, USA). Effect sizes (Hedges’ g) in absolute values were calculated to assess the strength of the intervention effects, were 0.2 ≤ g < 0.5 indicate small, 0.5 ≤ g < 0.8 indicate medium, and g ≥ 0.8 indicate large effects [61].

## Results

There were no significant differences in age (p = 0.421), body height (p = 0.361) and mass (p = 0.382) between intervention and control group. No baseline differences between groups were observed for the maximum plantar flexion moment (p = 0.894), AT stiffness (p = 0.421) and energetic cost (p = 0.143, tab. 1). Both the maximum plantar flexion moment and AT stiffness increased significantly in the intervention group (p = 0.024, p = 0.048) without any significant changes in the control group (p = 0.296, p = 0.745, tab. 1). Furthermore, we found a significant decrease in the energetic cost of running following the 14 weeks of training in the intervention group (p = 0.028) and no significant changes in the control group (p = 0.688, tab. 1). Both groups did not show any significant changes in the strike index (intervention p = 0.868, control p = 0.868), stance time (p = 0.283, p = 0.283), flight time (p = 0.981, p = 0.252) and cadence (p = 0.310, p = 0.384, tab. 1) after the 14 weeks of training, indicating that the training did not influence the foot strike pattern.

Following the intervention, ankle and knee joint kinematics did not significantly change during the stance phase of running, i.e. joint angles at touch down (ankle p = 0.108, knee p = 0.064), toe-off (p = 0.161, p = 0.844), maximal ankle dorsiflexion (p = 0.576), and maximal knee flexion (p = 0.138, tab. 2, fig. 2). The soleus MTU showed a lengthening-shortening behaviour during the stance phase, with shortening starting at 59 ± 2% of the stance phase similarly pre and post intervention (p = 0.266, g = 0.30, fig. 3). The training had no effect on the MTU length, length changes and velocity, neither when averaged over the entire stance phase (p = 0.943, p = 0.273, p = 0.274) nor over the subphase of MTU lengthening (p = 0.931, p = 0.893, p = 0.788) or MTU shortening (p = 0.946, p = 0.470, p = 0.189, tab. 3, fig. 3). Despite the MTU lengthening, the soleus muscle fascicles shortened continuously throughout the entire stance phase of running (fig. 3). Following the intervention, the fascicle shortening was not significantly different over the entire stance phase (p = 0.662) and the phase of MTU lengthening (p = 0.106) but in the phase of MTU shortening (p = 0.016, tab. 3). L_0_ (pre 43.1 ± 5.7 mm, post 44.1 ± 8.9 mm, p = 0.767, g = 0.08) and thus V_max_ (pre 291 ± 38 mm/s, post 298 ± 17 mm/s, p = 0.767, g = 0.08) was not significantly altered due to training. The operating fascicle length averaged over the stance phase (pre 0.87 ± 0.11 L_0_, post 0.85 ± 0.13 L_0_, p = 0.360, g = 0.16) but also during the phase of MTU lengthening (pre 0.92 ± 0.12 L_0_, post 0.91 ± 0.15 L_0_, p = 0.772, g = 0.07) and shortening (pre 0.81 ± 0.10 L_0_, post 0.76 ± 0.11 L_0_, p = 0.226, g = 0.32) was not significantly changed following training. Consequently, the force-length potential was not significantly different between pre and post training in the different phases (stance phase p = 0.172, g = 0.14, MTU lengthening p = 0.713, g = 0.10, MTU shortening p = 0.640, g = 0.12, fig. 4).

**Fig. 2:**
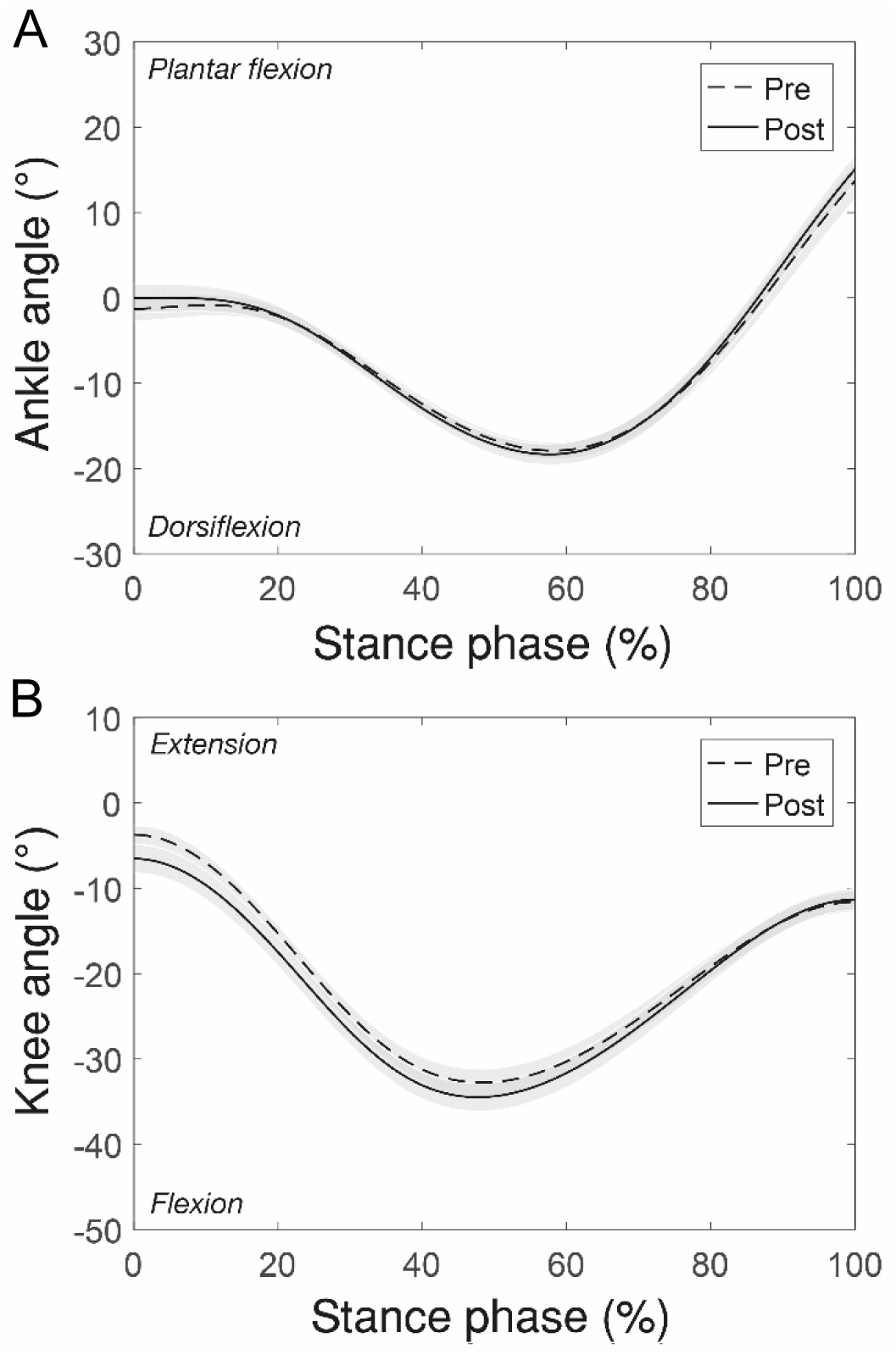
A) Ankle joint angle and B) knee joint angle during the stance phase of running before and after the training intervention (mean±SE, n=13).

**Fig. 3:**
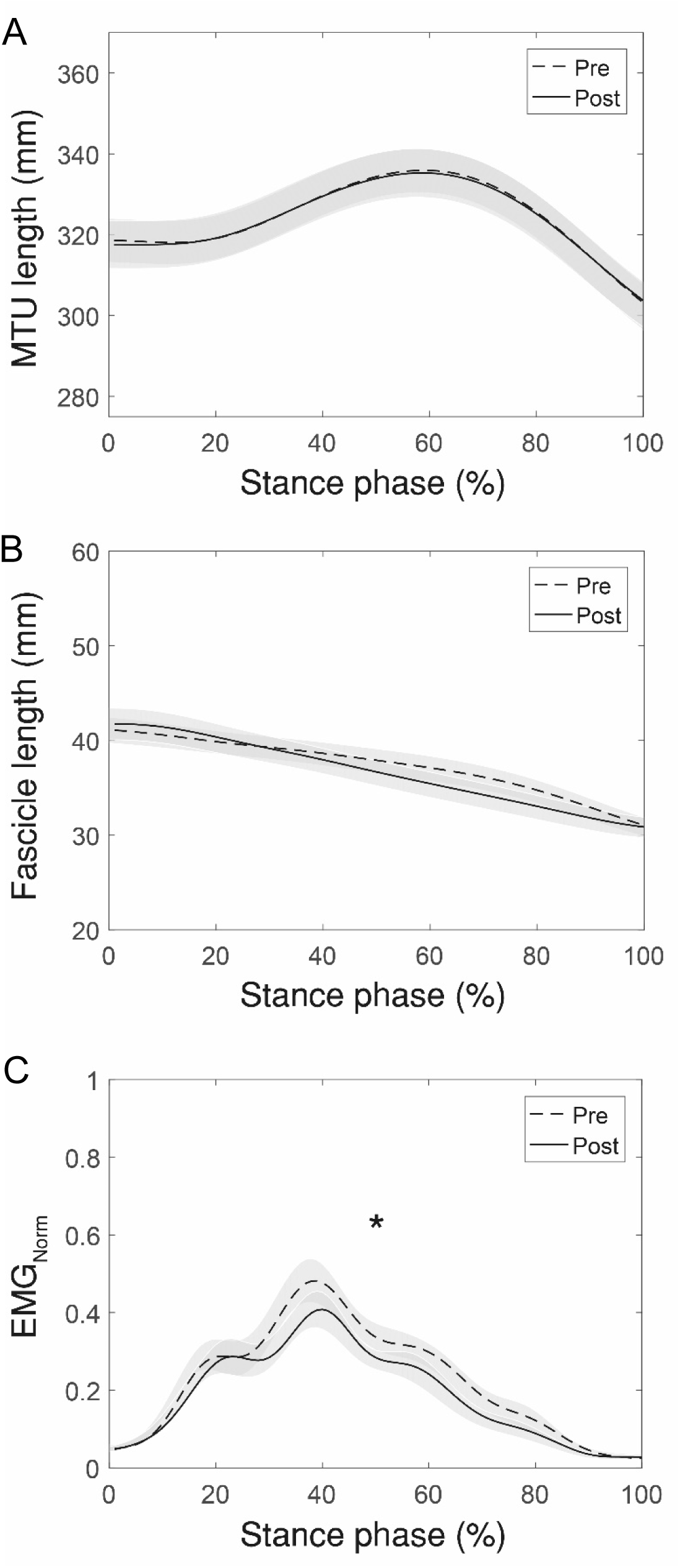
A) Soleus muscle-tendon unit (MTU) length, B) muscle fascicle length and C) electromyographic (EMG) activity (normalized to a maximum voluntary isometric contraction) during the stance phase of running before and after the training intervention (mean±SE, n=13). *Significant difference of the stance phase-averaged EMG activation between pre and post (p<0.05).

**Fig. 4:**
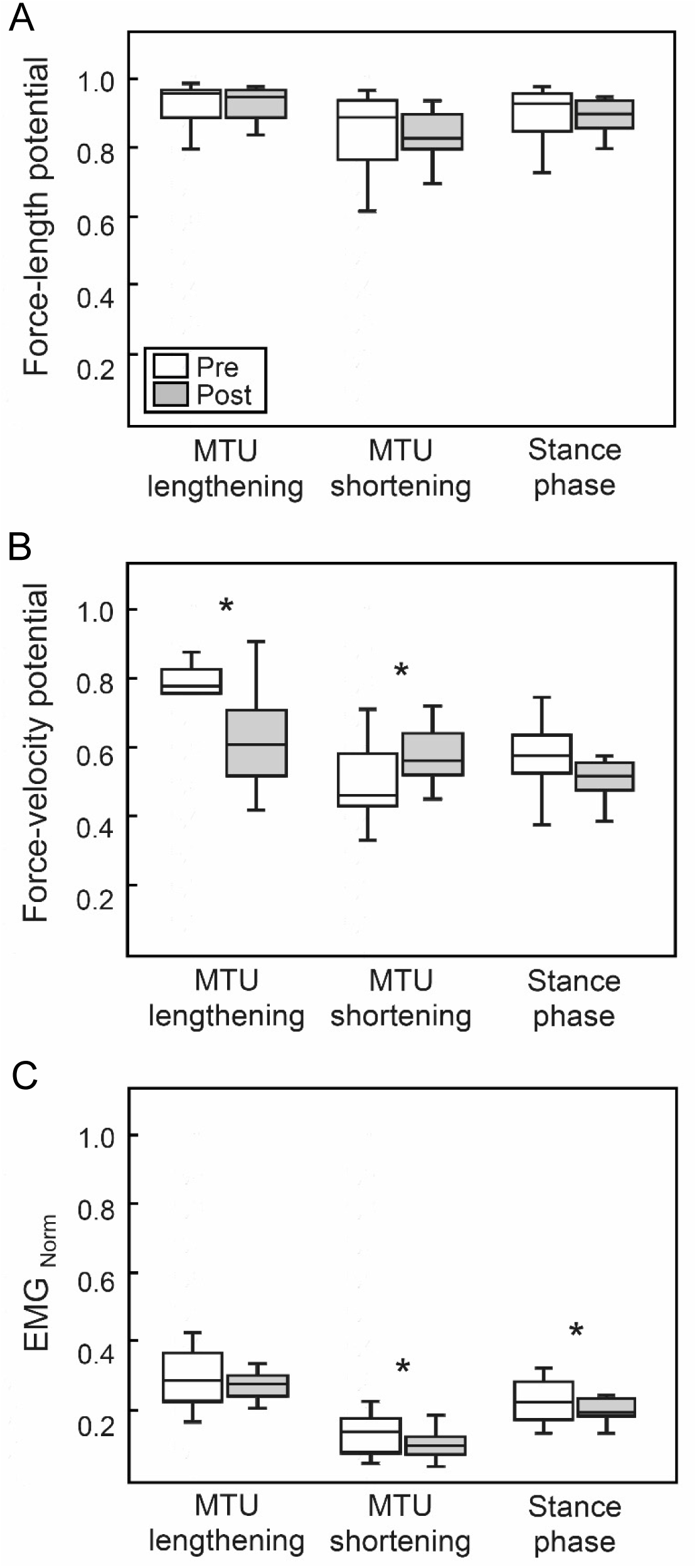
A) Soleus fascicle force-length potential, B) force-velocity potential and C) electromyographic (EMG) activity (normalized to a maximum voluntary isometric contraction), averaged over the phase of MTU lengthening, MTU shortening and the entire stance phase of running before and after the training intervention (n=13). *Significant difference between pre and post (p<0.05).

Following the training, the soleus force-velocity potential was significantly lower in the phase of MTU lengthening (p = 0.030, g = 0.64) and significantly higher when the MTU shortened (p = 0.045, g = 0.58) with no significant difference over the entire stance (p = 0.249, g = 0.31, fig. 4). This was the consequence of a tendency towards higher fascicle shortening velocity during MTU lenthgening (pre 0.088 ± 0.054 V_max_, post 0.129 ± 0.061 V_max_, p = 0.073, g = 0.51) and a significantly lower velocity during MTU shortening after training (pre 0.174 ± 0.057 V_max_, post 0.127 ± 0.008 V_max_, p = 0.007, g = 0.83. Furthermore, the averaged EMG activation over the phase of MTU shortening (p = 0.028, g = 0.67) and the entire stance phase of running was significantly reduced following the intervention (p = 0.017, g = 0.60, fig. 3 & fig. 4). Compared to pre-intervention running, the fascicle velocity in the phase of MTU lengthening was closer to the velocity for optimal enthalpy efficiency after the training (fig. 5). Consequently, the fascicles operated at a significantly higher enthalpy efficiency in the phase of MTU lengthening after the training (p = 0.006, g = 0.85, fig. 5 & fig. 6), while there was no significant pre-post difference in the phase of MTU shortening (p = 0.640, g = 0.12, fig. 6). Over the entire stance phase of running the enthalpy efficiency of the fascicle shortening was also significantly increased following the training (p = 0.025, g = 0.66, fig. 6).

**Fig. 5:**
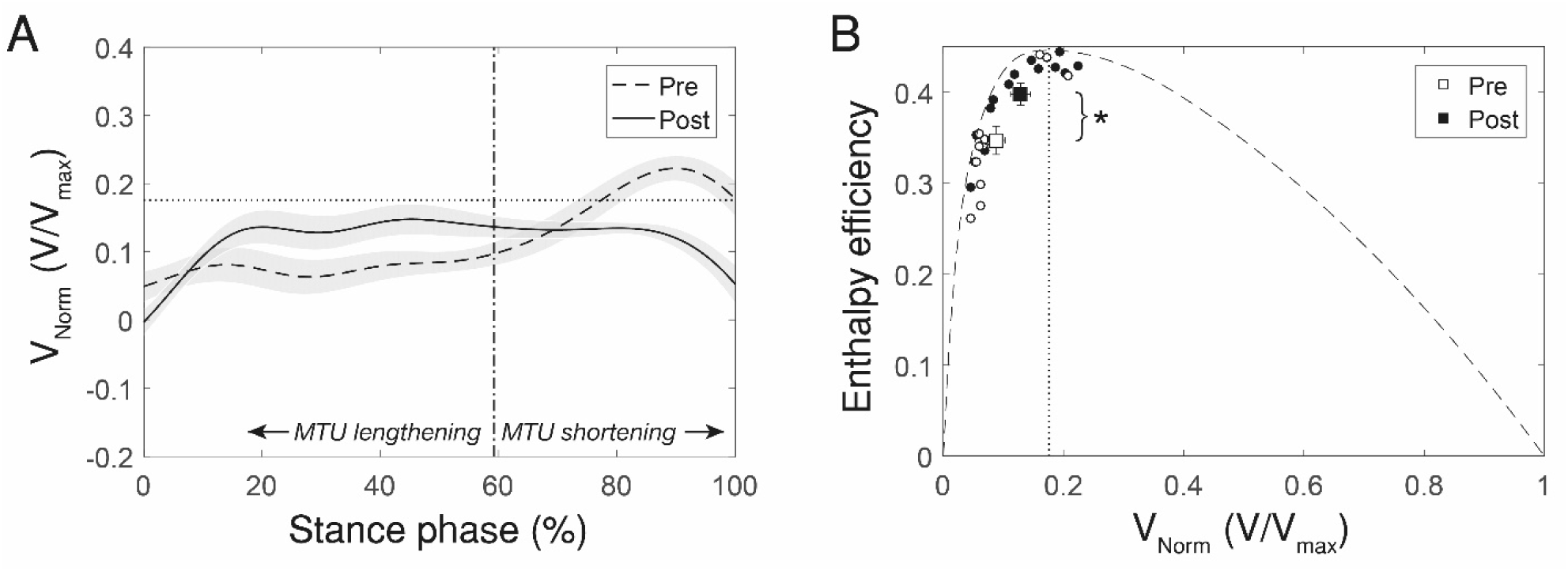
A) Soleus muscle fascicle operating velocity over the stance phase of running before and after the intervention (mean±SE) and velocity of maximum enthalpy efficiency (i.e. 0.18 V_max_, horizontal dashed line). Following the intervention, the fascicle shortening velocity was closer to the velocity optimal for maximum enthalpy efficiency during most of the muscle-tendon unit (MTU) lengthening phase. B) Enthalpy efficiency-fascicle velocity relationship with average values of the phase of MTU lengthening, showing that the fascicles operated at a significantly higher enthalpy efficiency following the intervention (*p<0.05). Circles indicate the single participant values before (white) and after (black) the intervention and squares show the respective mean with standard error bars (n=13). The vertical dotted line shows the velocity of maximum efficiency.

**Fig. 6:**
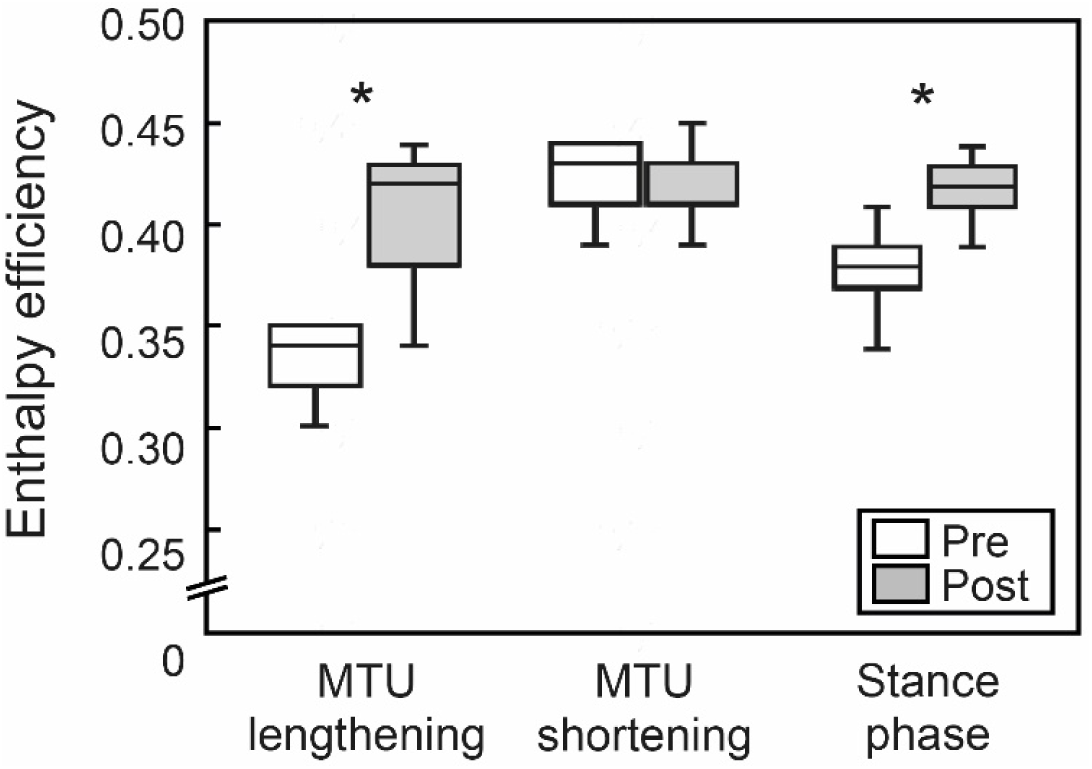
Soleus muscle fascicle enthalpy efficiency averaged over the phase of muscle-tendon unit (MTU) lengthening, MTU shortening and the entire stance phase of running before and after the training intervention (n=13). *Significant difference between pre and post (p<0.05).

## Discussion

Our current study showed for the first time that a specific muscle-tendon training that increases plantar flexor muscle strength and AT stiffness facilitates the enthalpy efficiency of the operating soleus muscle during the stance phase of running. The increased enthalpy efficiency was found in the first part of the stance phase where the soleus muscle produces work by active shortening and transfers muscular work to the tendon as strain energy. Furthermore, the results provide additional evidence that a combination of greater plantar flexor muscle strength and Achilles tendon stiffness decrease the energy cost of running [18,43] and strongly indicate that the soleus enthalpy efficiency is the explaining determinant. Following the intervention, the energetic cost of running was significantly reduced by about -4%. At the same time, the soleus, which is the main muscle for work/energy production during running [15,16], operated at a significantly increased (+7%) enthalpy efficiency throughout the stance phase. The enthalpy efficiency quantifies the portion of energy from ATP hydrolysis used by a muscle that is converted into mechanical muscular work [25]. Enthalpy efficiency depends on the velocity of muscle shortening with a steep increase at low velocities until the peak at around 0.18 V_max_ and again decreasing at higher shortening velocities [24,25]. For the whole stance phase, fascicle shortening, the force-length potential and the force-velocity potential of the soleus muscle was not significantly different before and after the intervention, indicating a similar energy production through muscular work of the soleus muscle. During the propulsion phase of running (i.e. MTU shortening), where both tendon and muscle transfer energy/work to the skeleton [62,63], the enthalpy efficiency of the operating soleus muscle was high both pre and post intervention (94% and 93% of the maximum efficiency). In contrast, during the first part of the stance phase (i.e. MTU lengthening), where energy is transferred from the contractile element to the tendon, the enthalpy efficiency was lower during pre-intervention running (77% of the maximum efficiency). The relevant part of the soleus fascicle shortening occurred during this first part of stance (59% of the entire shortening range). In combination with the high muscle activation (muscle activation was higher during MTU lengthening than during MTU shortening), this indicates an important energy production through muscular work during MTU lengthening.

The exercise-induced increase in muscle strength and AT stiffness resulted in an alteration of the operating fascicle velocity profile that led to a significant increase of the enthalpy efficiency of the operating soleus in the phase of MTU lengthening (88% of the maximum efficiency), improving the efficiency of muscular work production. The significant increase of the enthalpy efficiency following training in the phase of MTU lengthening demontrates that a substantial part of the entire muscular work was generated more economically. In the second part of the stance phase, where the MTU shortened, the high efficiency was maintained after the intervention and, further, the fascicles operated at a significantly higher force-velocity potential. This was possible due to a shift of the shortening velocity around the plateau of the efficiency-velocity curve, from the descending part of the plateau before the training to the ascending part after training (fig. 5), without a significant decline in the enthalpy efficiency. Consequently, the overall enthalpy efficiency throughout the stance phase of each step during running was increased. Furthermore, the higher force-velocity potential in the phase of MTU shortening due to training was accompanied by a reduced soleus EMG activation. The overall EMG activity during the stance phase was significantly lower as well after the intervention. This may indicate that a less active soleus muscle volume was required during running, which could – in addition to the higher operating enthalpy efficiency – reduce the metabolic energy cost [5,14,64].

Previous studies provided evidence that the cost of force to support the body mass and the time course of force application to the ground are the major determinants of the energetic cost of running [8,65,66]. According to the ‘cost of generating force hypothesis’ [8], the rate of metabolic energy consumption is directly related to the body mass and the time available to generate force, which results in a constant cost coefficient (i.e. energy required per unit force). However, modifications in the muscle effective mechanical advantage (i.e. ratio of the muscle moment arm to the moment arm of the ground reaction force [67]) within the lower extremities can influence the cost coefficient of human locomotion [68,69]. In our study, the metabolic cost of running was reduced after the training without any changes in the contact time and body mass, indicating a decrease of the cost coefficient. The similar strike index and lower leg kinematics before and after the intervention suggest unchanged effective mechanical advantages within the lower extremities and, therefore, this would not be the reason for the reduced cost coefficient. Instead, our findings show that an adjusted time course of the shortening velocity of the soleus muscle during the stance phase of running following the training can influence the cost coefficient as a result of increased enthalpy efficiency of the operating soleus and, thus, complement the earlier studies on the mechanical advantage and cost coefficient interaction [66,67]. The observed continuous soleus fascicle shortening behavior during the stance phase of running in our study is in agreement with other in vivo experiments using the ultrasound methodology and comparable running speeds [23,70–72]. The importance of the energy production by the plantar flexor muscles for the propulsion phase (i.e. shortening of the MTU) during running is well accepted [23,73–75] because the mechanical power produced at the ankle joint in this phase is highest and determines running performance [76–78]. Our current results regarding the enthalpy efficiency of muscular energy gain and running economy show for the first time that also the phase of the MTU lengthening is crucial for the overall metabolic energy consumption during running.

The findings of the current study provide further evidence [20,43] that a strength training of the plantar flexors has the potential to improve running economy. We used a specific high intensity muscle-tendon training [28,29,33], targeting an adaptation of both AT stiffness and plantar flexor muscle strength [18,43], to maintain the functional integrity of the contractile and series-elastic element [79]. Strength increases without concomitant stiffening of the AT after a period of training may increase levels of operating and maximum AT strain [29,80], which have been associated with pathologies [81] but also possible functional decline [82,83]. In our study, the maximum AT strain during the MVCs was not affected by the training (pre 6.2 ± 1.6%, post 6.0 ± 1.2%, p = 0.501) despite an increase in the muscle strength of the plantar flexors, indicating a balanced adaptation of muscle and tendon. Therefore, a specific muscle and tendon training [28,29,33] can be recommended to increase endurance performance and a specific diagnostic of muscle and tendon properties may allow tailoring the training to individual deficits of muscle strength or tendon stiffness [79].

To assess the enthalpy efficiency-shortening velocity relationship, we used a biologically founded value of V_max_, i.e. 6.77 L_0_ /s. However, during submaximal running, the lower activation level and selective slow fibre type recruitment may affect the actual enthalpy efficiency-shortening velocity relationship. Furthermore, differences in fibre type distribution may also affect the shape of the enthalpy efficiency-shortening velocity curve [26]. We evaluated the effect of a) decreasing and increasing V_max_ by 10% intervals and b) replacing the underlying enthalpy efficiency values from Hill (1964) [24] by the data presented by Barclay et al. (1993) [26] for the predominantly slow fiber type soleus mouse muscle, comparable to the human soleus muscle. The significant pre to post enthalpy efficiency increase for the MTU lengthening phase and entire stance phase persisted for values between V_max-30%_ and V_max+10%_ both using the data of Hill or Barclay and colleagues (p<0.05), which confirms and strengthens the observed intervention effect.

In conclusion, the current study gives new insights into the soleus muscle mechanics and metabolic energetics during human running. In support of our earlier study, an exercised-induced increase of plantar flexor muscle strength and AT stiffness reduced the metabolic energy cost of running. We found the reason for this improvement to be an alteration in the soleus fascicle velocity profile throughout the stance phase, which led to a significantly higher enthalpy efficiency of the operating soleus muscle. The enthalpy efficiency was particularly increased in the phase of muscle-tendon unit lengthening, where the activation is high and the soleus generates an important part of the mechanical energy required for running.

## Authors’ contributions

S.B., F.M., A.S. and A.A. designed research; S.B., F.M. and A.S. performed research; S.B. analyzed data; S.B. and A.A. drafted the manuscript and F.M. and A.S. made important intellectual contributions during revision.

## Acknowledgments

We acknowledge the support of Antonis Ekizos, Arno Schroll, Leon Brüll and Victor Munoz-Martel for data recording and analysis.

## References

1. Pontzer H. 2017 Economy and Endurance in Human Evolution. Curr. Biol. 27, R613–R621. (doi: 10.1016/j.cub.2017.05.031)

2. Bramble DM, Lieberman DE. 2004 Endurance running and the evolution of Homo. Nature 432, 345–352. (doi: 10.1038/nature03052)

3. Raichlen DA, Polk JD. 2013 Linking brains and brawn: exercise and the evolution of human neurobiology. Proc. R. Soc. B Biol. Sci. 280, 20122250. (doi: 10.1098/rspb.2012.2250)

4. Carrier DR, Kapoor AK, Kimura T, Nickels MK, Satwanti, Scott EC, So JK, Trinkaus E. 1984 The Energetic Paradox of Human Running and Hominid Evolution. Curr. Anthropol. 25, 483–495.

5. Fletcher JR, MacIntosh BR. 2017 Running Economy from a Muscle Energetics Perspective. Front. Physiol. 8, 433. (doi: 10.3389/fphys.2017.00433)

6. Kipp S, Byrnes WC, Kram R. 2018 Calculating metabolic energy expenditure across a wide range of exercise intensities: the equation matters. Appl. Physiol. Nutr. Metab. Physiol. Appl. Nutr. Metab. 43, 639–642. (doi: 10.1139/apnm-2017-0781)

7. Joyner MJ. 1991 Modeling: optimal marathon performance on the basis of physiological factors. J. Appl. Physiol. 70, 683–687. (doi: 10.1152/jappl.1991.70.2.683)

8. Kram R, Taylor CR. 1990 Energetics of running: a new perspective. Nature 346, 265–267. (doi: 10.1038/346265a0)

9. Arellano CJ, Kram R. 2014 Partitioning the Metabolic Cost of Human Running: A Task-by-Task Approach. Integr. Comp. Biol. 54, 1084–1098. (doi: 10.1093/icb/icu033)

10. Gordon AM, Huxley AF, Julian FJ. 1966 The variation in isometric tension with sarcomere length in vertebrate muscle fibres. J. Physiol. 184, 170–192.

11. Hill Archibald Vivian. 1938 The heat of shortening and the dynamic constants of muscle. Proc. R. Soc. Lond. Ser. B - Biol. Sci. 126, 136–195. (doi: 10.1098/rspb.1938.0050)

12. Bohm S, Mersmann F, Santuz A, Arampatzis A. 2019 The force–length–velocity potential of the human soleus muscle is related to the energetic cost of running. Proc. R. Soc. B Biol. Sci. 286, 20192560. (doi: 10.1098/rspb.2019.2560)

13. Bohm S, Marzilger R, Mersmann F, Santuz A, Arampatzis A. 2018 Operating length and velocity of human vastus lateralis muscle during walking and running. Sci. Rep. 8, 5066. (doi: 10.1038/s41598-018-23376-5)

14. Roberts TJ, Marsh RL, Weyand PG, Taylor CR. 1997 Muscular Force in Running Turkeys: The Economy of Minimizing Work. Science 275, 1113–1115. (doi: 10.1126/science.275.5303.1113)

15. Hamner SR, Delp SL. 2013 Muscle contributions to fore-aft and vertical body mass center accelerations over a range of running speeds. J. Biomech. 46, 780–787. (doi: 10.1016/j.jbiomech.2012.11.024)

16. Dorn TW, Schache AG, Pandy MG. 2012 Muscular strategy shift in human running: dependence of running speed on hip and ankle muscle performance. J. Exp. Biol. 215, 1944–1956. (doi: 10.1242/jeb.064527)

17. Fletcher JR, MacIntosh BR. 2015 Achilles tendon strain energy in distance running: consider the muscle energy cost. J. Appl. Physiol. 118, 193–199. (doi: 10.1152/japplphysiol.00732.2014)

18. Arampatzis A, De Monte G, Karamanidis K, Morey-Klapsing G, Stafilidis S, Brueggemann G-P. 2006 Influence of the muscle-tendon unit’s mechanical and morphological properties on running economy. J. Exp. Biol. 209, 3345–3357. (doi: 10.1242/jeb.02340)

19. Albracht K, Arampatzis A. 2013 Exercise-induced changes in triceps surae tendon stiffness and muscle strength affect running economy in humans. Eur. J. Appl. Physiol. 113, 1605–1615. (doi: 10.1007/s00421-012-2585-4)

20. Fletcher JR, Esau SP, MacIntosh BR. 2010 Changes in tendon stiffness and running economy in highly trained distance runners. Eur. J. Appl. Physiol. 110, 1037–1046. (doi: 10.1007/s00421-010-1582-8)

21. Rogers SA, Whatman CS, Pearson SN, Kilding AE. 2017 Assessments of Mechanical Stiffness and Relationships to Performance Determinants in Middle-Distance Runners. Int. J. Sports Physiol. Perform. 12, 1329–1334. (doi: 10.1123/ijspp.2016-0594)

22. Albracht K, Arampatzis A, Baltzopoulos V. 2008 Assessment of muscle volume and physiological cross-sectional area of the human triceps surae muscle in vivo. J. Biomech. 41, 2211–2218. (doi: 10.1016/j.jbiomech.2008.04.020)

23. Lai A, Lichtwark GA, Schache AG, Lin Y-C, Brown NAT, Pandy MG. 2015 In vivo behavior of the human soleus muscle with increasing walking and running speeds. J. Appl. Physiol. Bethesda Md 1985 118, 1266–1275. (doi: 10.1152/japplphysiol.00128.2015)

24. Hill AV. 1964 The efficiency of mechanical power development during muscular shortening and its relation to load. Proc. R. Soc. Lond. B Biol. Sci. 159, 319–324. (doi: 10.1098/rspb.1964.0005)

25. Barclay CJ. 2015 Energetics of contraction. Compr. Physiol. 5, 961–995. (doi: 10.1002/cphy.c140038)

26. Barclay CJ, Constable JK, Gibbs CL. 1993 Energetics of fast- and slow-twitch muscles of the mouse. J. Physiol. 472, 61–80. (doi: 10.1113/jphysiol.1993.sp019937)

27. Folland DJP, Williams AG. 2007 Morphological and Neurological Contributions to Increased Strength. Sports Med. 37, 145–168. (doi: 10.2165/00007256-200737020-00004)

28. Bohm S, Mersmann F, Arampatzis A. 2015 Human tendon adaptation in response to mechanical loading: a systematic review and meta-analysis of exercise intervention studies on healthy adults. Sports Med. - Open 1, 7. (doi: 10.1186/s40798-015-0009-9)

29. Arampatzis A, Karamanidis K, Albracht K. 2007 Adaptational responses of the human Achilles tendon by modulation of the applied cyclic strain magnitude. J. Exp. Biol. 210, 2743–2753. (doi: 10.1242/jeb.003814)

30. Faul F, Erdfelder E, Lang A-G, Buchner A. 2007 G*Power 3: A flexible statistical power analysis program for the social, behavioral, and biomedical sciences. Behav. Res. Methods 39, 175–191. (doi: 10.3758/BF03193146)

31. Cavanagh PR, Lafortune MA. 1980 Ground reaction forces in distance running. J. Biomech. 13, 397–406. (doi: 10.1016/0021-9290(80)90033-0)

32. Nikolaidou ME, Marzilger R, Bohm S, Mersmann F, Arampatzis A. 2017 Operating length and velocity of human M. vastus lateralis fascicles during vertical jumping. R. Soc. Open Sci. 4, 170185. (doi: 10.1098/rsos.170185)

33. Bohm S, Mersmann F, Tettke M, Kraft M, Arampatzis A. 2014 Human achilles tendon plasticity in response to cyclic strain: effect of rate and duration. J. Exp. Biol. 217, 4010–4017. (doi: 10.1242/jeb.112268)

34. Arampatzis A, Morey-Klapsing G, Karamanidis K, DeMonte G, Stafilidis S, Brüggemann G-P. 2005 Differences between measured and resultant joint moments during isometric contractions at the ankle joint. J. Biomech. 38, 885–892. (doi: 10.1016/j.jbiomech.2004.04.027)

35. Arampatzis A, Karamanidis K, De Monte G, Stafilidis S, Morey-Klapsing G, Brüggemann G-P. 2004 Differences between measured and resultant joint moments during voluntary and artificially elicited isometric knee extension contractions. Clin. Biomech. 19, 277–283. (doi: 10.1016/j.clinbiomech.2003.11.011)

36. Mademli L, Arampatzis A, Morey-Klapsing G, Brüggemann G-P. 2004 Effect of ankle joint position and electrode placement on the estimation of the antagonistic moment during maximal plantarflexion. J. Electromyogr. Kinesiol. 14, 591–597. (doi: 10.1016/j.jelekin.2004.03.006)

37. An KN, Takahashi K, Harrigan TP, Chao EY. 1984 Determination of muscle orientations and moment arms. J. Biomech. Eng. 106, 280–282.

38. Fath F, Blazevich AJ, Waugh CM, Miller SC, Korff T. 2010 Direct comparison of in vivo Achilles tendon moment arms obtained from ultrasound and MR scans. J. Appl. Physiol. Bethesda Md 1985 109, 1644–1652. (doi: 10.1152/japplphysiol.00656.2010)

39. Maganaris CN, Baltzopoulos V, Sargeant AJ. 1998 Changes in Achilles tendon moment arm from rest to maximum isometric plantarflexion: in vivo observations in man. J. Physiol. 510, 977–985. (doi: 10.1111/j.1469-7793.1998.977bj.x)

40. Arampatzis A, Monte GD, Karamanidis K. 2008 Effect of joint rotation correction when measuring elongation of the gastrocnemius medialis tendon and aponeurosis. J. Electromyogr. Kinesiol. Off. J. Int. Soc. Electrophysiol. Kinesiol. 18, 503–508. (doi: 10.1016/j.jelekin.2006.12.002)

41. Schulze F, Mersmann F, Bohm S, Arampatzis A. 2012 A wide number of trials is required to achieve acceptable reliability for measurement patellar tendon elongation in vivo. Gait Posture 35, 334–338. (doi: 10.1016/j.gaitpost.2011.09.107)

42. De Monte G, Arampatzis A, Stogiannari C, Karamanidis K. 2006 In vivo motion transmission in the inactive gastrocnemius medialis muscle–tendon unit during ankle and knee joint rotation. J. Electromyogr. Kinesiol. 16, 413–422. (doi: 10.1016/j.jelekin.2005.10.001)

43. Albracht K, Arampatzis A. 2013 Exercise-induced changes in triceps surae tendon stiffness and muscle strength affect running economy in humans. Eur. J. Appl. Physiol. 113, 1605–1615. (doi: 10.1007/s00421-012-2585-4)

44. Péronnet F, Massicotte D. 1991 Table of nonprotein respiratory quotient: an update. Can. J. Sport Sci. 16, 23–9.

45. Saunders PU, Pyne DB, Telford RD, Hawley JA. 2004 Factors Affecting Running Economy in Trained Distance Runners. Sports Med. 34, 465–485. (doi: 10.2165/00007256-200434070-00005)

46. Fellin RE, Rose WC, Royer TD, Davis IS. 2010 Comparison of methods for kinematic identification of footstrike and toe-off during overground and treadmill running. J. Sci. Med. Sport 13, 646–650. (doi: 10.1016/j.jsams.2010.03.006)

47. Santuz A, Ekizos A, Arampatzis A. 2016 A Pressure Plate-Based Method for the Automatic Assessment of Foot Strike Patterns During Running. Ann. Biomed. Eng. 44, 1646–1655. (doi: 10.1007/s10439-015-1484-3)

48. Lutz GJ, Rome LC. 1996 Muscle function during jumping in frogs. I. Sarcomere length change, EMG pattern, and jumping performance. Am. J. Physiol. 271, C563–570.

49. Hawkins D, Hull ML. 1990 A method for determining lower extremity muscle-tendon lengths during flexion/extension movements. J. Biomech. 23, 487–494. (doi: 10.1016/0021-9290(90)90304-L)

50. Marzilger R, Legerlotz K, Panteli C, Bohm S, Arampatzis A. 2018 Reliability of a semi-automated algorithm for the vastus lateralis muscle architecture measurement based on ultrasound images. Eur. J. Appl. Physiol. 118, 291–301. (doi: 10.1007/s00421-017-3769-8)

51. Giannakou E, Aggeloussis N, Arampatzis A. 2011 Reproducibility of gastrocnemius medialis muscle architecture during treadmill running. J. Electromyogr. Kinesiol. Off. J. Int. Soc. Electrophysiol. Kinesiol. 21, 1081–1086. (doi: 10.1016/j.jelekin.2011.06.004)

52. Hof AL, van den Berg Jw. 1977 Linearity between the weighted sum of the EMGs of the human triceps surae and the total torque. J. Biomech. 10, 529–539. (doi: 10.1016/0021-9290(77)90033-1)

53. Luden N, Minchev K, Hayes E, Louis E, Trappe T, Trappe S. 2008 Human vastus lateralis and soleus muscles display divergent cellular contractile properties. Am. J. Physiol. Regul. Integr. Comp. Physiol. 295, R1593–1598. (doi: 10.1152/ajpregu.90564.2008)

54. Ranatunga KW. 1984 The force-velocity relation of rat fast- and slow-twitch muscles examined at different temperatures. J. Physiol. 351, 517–529. (doi: 10.1113/jphysiol.1984.sp015260)

55. Edgerton VR, Smith JL, Simpson DR. 1975 Muscle fibre type populations of human leg muscles. Histochem. J. 7, 259–266.

56. Larsson L, Moss RL. 1993 Maximum velocity of shortening in relation to myosin isoform composition in single fibres from human skeletal muscles. J. Physiol. 472, 595–614. (doi: 10.1113/jphysiol.1993.sp019964)

57. Johnson MA, Polgar J, Weightman D, Appleton D. 1973 Data on the distribution of fibre types in thirty-six human muscles. An autopsy study. J. Neurol. Sci. 18, 111–129. (doi: 10.1016/0022-510x(73)90023-3)

58. Winters JM, Stark L. 1985 Analysis of Fundamental Human Movement Patterns Through the Use of In-Depth Antagonistic Muscle Models. IEEE Trans. Biomed. Eng. BME-32, 826–839. (doi: 10.1109/TBME.1985.325498)

59. Winters JM, Stark L. 1988 Estimated mechanical properties of synergistic muscles involved in movements of a variety of human joints. J. Biomech. 21, 1027–1041. (doi: 10.1016/0021-9290(88)90249-7)

60. Umberger BR, Gerritsen KGM, Martin PE. 2003 A Model of Human Muscle Energy Expenditure. Comput. Methods Biomech. Biomed. Engin. 6, 99–111. (doi: 10.1080/1025584031000091678)

61. Cohen J. 1988 Statistical Power Analysis for the Behavioral Sciences. Psychology Press.

62. Farris DJ, Sawicki GS. 2012 Human medial gastrocnemius force–velocity behavior shifts with locomotion speed and gait. Proc. Natl. Acad. Sci. 109, 977–982. (doi: 10.1073/pnas.1107972109)

63. Lai A, Schache AG, Lin Y-C, Pandy MG. 2014 Tendon elastic strain energy in the human ankle plantar-flexors and its role with increased running speed. J. Exp. Biol. 217, 3159–3168. (doi: 10.1242/jeb.100826)

64. Roberts TJ, Azizi E. 2011 Flexible mechanisms: the diverse roles of biological springs in vertebrate movement. J. Exp. Biol. 214, 353–361. (doi: 10.1242/jeb.038588)

65. Taylor CR, Heglund NC, McMahon TA, Looney TR. 1980 Energetic Cost of Generating Muscular Force During Running: A Comparison of Large and Small Animals. J. Exp. Biol. 86, 9–18.

66. Roberts TJ, Kram R, Weyand PG, Taylor CR. 1998 Energetics of bipedal running. I. Metabolic cost of generating force. J. Exp. Biol. 201, 2745–2751.

67. Biewener AA. 1998 Muscle Function in vivo: A Comparison of Muscles used for Elastic Energy Savings versus Muscles Used to Generate Mechanical Power1. Integr. Comp. Biol. 38, 703–717. (doi: 10.1093/icb/38.4.703)

68. Ekizos A, Santuz A, Arampatzis A. 2018 Short- and long-term effects of altered point of ground reaction force application on human running energetics. J. Exp. Biol., jeb.176719. (doi: 10.1242/jeb.176719)

69. Biewener AA, Farley CT, Roberts TJ, Temaner M. 2004 Muscle mechanical advantage of human walking and running: implications for energy cost. J. Appl. Physiol. 97, 2266–2274. (doi: 10.1152/japplphysiol.00003.2004)

70. Cronin NJ, Finni T. 2013 Treadmill versus overground and barefoot versus shod comparisons of triceps surae fascicle behaviour in human walking and running. Gait Posture 38, 528–533. (doi: 10.1016/j.gaitpost.2013.01.027)

71. Werkhausen A, Cronin NJ, Albracht K, Paulsen G, Larsen AV, Bojsen-Møller J, Seynnes OR. 2019 Training-induced increase in Achilles tendon stiffness affects tendon strain pattern during running. PeerJ 7. (doi: 10.7717/peerj.6764)

72. Rubenson J, Pires NJ, Loi HO, Pinniger GJ, Shannon DG. 2012 On the ascent: the soleus operating length is conserved to the ascending limb of the force–length curve across gait mechanics in humans. J. Exp. Biol. 215, 3539–3551. (doi: 10.1242/jeb.070466)

73. Stefanyshyn DJ, Nigg BM. 1998 Contribution of the lower extremity joints to mechanical energy in running vertical jumps and running long jumps. J. Sports Sci. 16, 177–186. (doi: 10.1080/026404198366885)

74. Komi PV. 2000 Stretch-shortening cycle: a powerful model to study normal and fatigued muscle. J. Biomech. 33, 1197–1206. (doi: 10.1016/S0021-9290(00)00064-6)

75. Novacheck TF. 1998 The biomechanics of running. Gait Posture 7, 77–95. (doi: 10.1016/S0966-6362(97)00038-6)

76. Buczek FL, Cavanagh PR. 1990 Stance phase knee and ankle kinematics and kinetics during level and downhill running. Med. Sci. Sports Exerc. 22, 669–677.

77. Arampatzis A, Bruggemann GP, Metzler V. 1999 The effect of speed on leg stiffness and joint kinetics in human running. J. Biomech. 32, 1349–1353. (doi: 10.1016/S0021-9290(99)00133-5)

78. Schache AG, Blanch PD, Dorn TW, Brown N a. T, Rosemond D, Pandy MG. 2011 Effect of Running Speed on Lower Limb Joint Kinetics. Med. Sci. Sports Exerc. 43, 1260–1271. (doi: 10.1249/MSS.0b013e3182084929)

79. Arampatzis A, Mersmann F, Bohm S. 2020 Individualized Muscle-Tendon Assessment and Training. Front. Physiol. 11. (doi: 10.3389/fphys.2020.00723)

80. Mersmann F, Bohm S, Arampatzis A. 2017 Imbalances in the Development of Muscle and Tendon as Risk Factor for Tendinopathies in Youth Athletes: A Review of Current Evidence and Concepts of Prevention. Front. Physiol. 8. (doi: 10.3389/fphys.2017.00987)

81. Obst SJ, Heales LJ, Schrader BL, Davis SA, Dodd KA, Holzberger CJ, Beavis LB, Barrett RS. 2018 Are the Mechanical or Material Properties of the Achilles and Patellar Tendons Altered in Tendinopathy? A Systematic Review with Meta-analysis. Sports Med. 48, 2179–2198. (doi: 10.1007/s40279-018-0956-7)

82. Lichtwark GA, Wilson AM. 2007 Is Achilles tendon compliance optimised for maximum muscle efficiency during locomotion? J. Biomech. 40, 1768–1775. (doi: 10.1016/j.jbiomech.2006.07.025)

83. Orselli MIV, Franz JR, Thelen DG. 2017 The effects of Achilles tendon compliance on triceps surae mechanics and energetics in walking. J. Biomech. 60, 227–231. (doi: 10.1016/j.jbiomech.2017.06.022)

